# Biomechanics of Finger Pad Response under Torsion

**DOI:** 10.1101/2022.11.07.515186

**Authors:** Sophie du Bois de Dunilac, David Córdova Bulens, Philippe Lefèvre, Stephen J. Redmond, Benoit P. Delhaye

## Abstract

Surface skin deformation of the finger pad during partial slippage at finger-object interfaces elicits tactile feedback. During object manipulation, torque is often present, which can cause partial slippage. Until now, studies of surface skin deformation have used stimuli sliding on rectilinear tangential trajectories.

Here we studied surface skin dynamics under torsion. A custom robotic platform stimulated the finger pad with a flat transparent surface, controlling the normal forces and rotation speeds applied while monitoring the contact interface using optical imaging.

We observed the characteristic pattern by which partial slips develop, starting at the periphery of the contact and propagating towards its centre, and the resulting surface strains. The 20-fold range of normal forces and angular velocities used highlights the effect of those parameters on the resulting torque and skin strains. While normal force increases the contact area, generated torque, strains, and twist angle required to reach full slip, angular velocity increases loss of contact at the periphery and strain rates (although not total strains). We also discuss the surprisingly large inter-individual variability in skin biomechanics, notably observed in the twist angle the stimulus needed to rotate before reaching full slip.

## Introduction

Humans are incredibly skilled at manipulating objects. Our success at dexterous manipulation relies on the feedback from our sensory systems, in particular tactile, proprioceptive, and visual. Amongst them, the contribution of tactile feedback is essential, in that removing it by trauma or anaesthesia of the fingers leads to major performance degradation even in the simplest manipulation tasks (Augurelle et al. 2003; Witney et al. 2004). Moreover, cutaneous feedback influences weight perception, and subsequently the grip force used during object manipulation (Park et al. 2020).

What we sense by touch is mediated by the mechanoreceptors, the sensory endings of the afferent nerves densely innervating the skin (Corniani and Saal 2020), which transduce mechanical deformations at the level of their end-organs into neural activity (Handler and Ginty 2021). Indeed, mechanical events that fail to induce sufficient deformation of the skin cannot be sensed (Barrea et al. 2018). Knowing how skin deforms under loading is a crucial step in understanding how our sense of touch is generated and subsequently used to guide our actions in tasks such as object manipulation (Johansson and Flanagan 2009).

While mechanoreceptors transduce local deformation of the skin, it is the integration of the information coming from numerous tactile fibers over a larger area of skin that enables us to decode the properties of the interface between the skin and the object being manipulated, such as normal and shear forces, slip, texture, friction, and torque, for example (Khamis, Birznieks, and Redmond 2015; Delhaye, Xia, and Bensmaia 2019; Weber et al. 2013; Cadoret and Smith 1996; Loutit et al. 2022). Through further inference, the properties of the object itself may be estimated; for example, its mass. To better understand the nature of this population-level tactile information, it is therefore relevant to study how the skin of the finger pad deforms when subjected to mechanical stimuli (Saal et al. 2017; Birznieks et al. 2009).

The contact interface between a passive finger (i.e., the participant is relaxed and does not attempt to move the finger) and a moving glass plate has been studied previously. When rectilinear tangential loads are applied to the finger pad, the periphery of the contact area between the skin and the glass plate starts slipping first. This slip annulus then propagates towards the centre of the contact area until all skin in contact with the glass plate is slipping (André et al. 2011). Surface strains are present at the border between the stuck area and the slip annulus (Delhaye et al. 2016), and can be linked with FA-I firing rates (Delhaye et al. 2021). The partial slip phase of the episode is of particular interest, as it might give us an interval of time during which we can sense the impending full slip and so adjust our grip force before the object in our hand starts to escape our grasp (Schiltz et al. 2021).

Most studies looking at the evolution of the slip annulus have focused on rectilinear motion (Lévesque and Hayward 2003; Tada and Kanade 2004; André et al. 2011; Delhaye, Lefèvre, and Thonnard 2014; Delhaye et al. 2016; Delhaye et al. 2021). However, in the likely event that the lifting force vector applied by the fingers to the object does not pass through the object’s centre of mass, the object will experience a torque and so a tendency to rotate. This torque can be counteracted by a reactive force arising from the friction at the finger-object interface, which can generate the required reactive torques to counter the object’s tendency to rotate (Kinoshita et al. 1997); for a two-finger precision grasp there are of course two finger-object interfaces (Goodwin, Jenmalm, and Johansson 1998), but the idea remains the same. The minimum grip force required to prevent slip arises from the resultant combination of tangential forces and torques (Kinoshita et al. 1997; Goodwin, Jenmalm, and Johansson 1998). In addition, the minimum grip force depends on the frictional condition (Kinoshita et al. 1997) and the curvature of the grasped surface (Goodwin, Jenmalm, and Johansson 1998), in response to which participants can appropriately scale their grip force. Cutaneous feedback elicited by torsion contains information about the applied torque. Passive experiments in monkey (Birznieks et al. 2010) and human fingers (Loutit et al. 2022) showed that the magnitude and direction of the torque applied by a flat stimulus could be extracted from the population response of sensory afferents, despite their complex responses at an individual level. Based on the link established between surface skin strains and neural recordings in rectilinear motion (Delhaye et al. 2021), we expect that some of the inter-afferent variability observed under torsion might be explained by specific local strains present in their receptive fields. Overall, understanding how skin deforms locally under torsion could help uncover unknown properties of the mechanotransduction process, and in a wider perspective, how humans scale their grip force in the presence of torques.

To date, and to our knowledge, there have been no studies on the mechanics of partial slip on the finger pad arising from torsion. This study is a first step toward understanding cutaneous feedback in the presence of torque. Here, we extend the work studying the dynamics of the finger pad under translation (André, Lefèvre, and Thonnard 2009; Delhaye, Lefèvre, and Thonnard 2014; Delhaye et al. 2016) to rotations. To that end, we passively stimulated the finger pad with a flat transparent surface, under controlled normal forces and rotation speeds, both of which were varied to span a range relevant to human touch in everyday scenarios, while monitoring the contact interface using optical imaging. We report the characteristic pattern by which the slip develops, the strong influence of force and speed, and discuss the large inter-individual variability that suggests exercising caution when analysing data coming from multiple participants.

## Methods

### Participants

Seven healthy human volunteers (aged 23-35, 4 males) participated in the study. All provided informed consent before participating in this study. The experimental procedure was approved by the local ethics committee at the Université catholique de Louvain.

### Experimental protocol

A platform consisting of an industrial four-axis robot (DENSO HS4535G), two six-axis force sensors (ATI IA, Mini 40), a glass plate, a finger holder, and an imaging system was used to perform the experiment (Fig. 1A). More details about the setup can be found in Delhaye *et al*. (Delhaye, Lefèvre, and Thonnard 2014; Delhaye et al. 2016). During a trial, the plate was brought into contact with the participant’s finger until the desired normal force level (*F*_*N*_) was reached. Once the normal force was stable, the robot rotated the plate at a constant angular velocity (ω) until a rotation of 80° was reached, and then stayed at that position for 1 s. The plate position was adjusted before the experiment such that the centre of rotation was as close as possible to the centre of the contact area. Five different normal force levels were tested (0.5, 1, 2, 5, 10 N) at an angular velocity of 20 °/s, and 5 different angular velocities were tested (5, 10, 20, 50, 100 °/s) at a normal force of 2 N (Fig. 1C), thus a 20-fold range for both parameters. Each condition was repeated 5 times in clockwise and counterclockwise directions, for a total of 90 trials. The same glass plate was used for all participants and trials.

**Figure 1.**
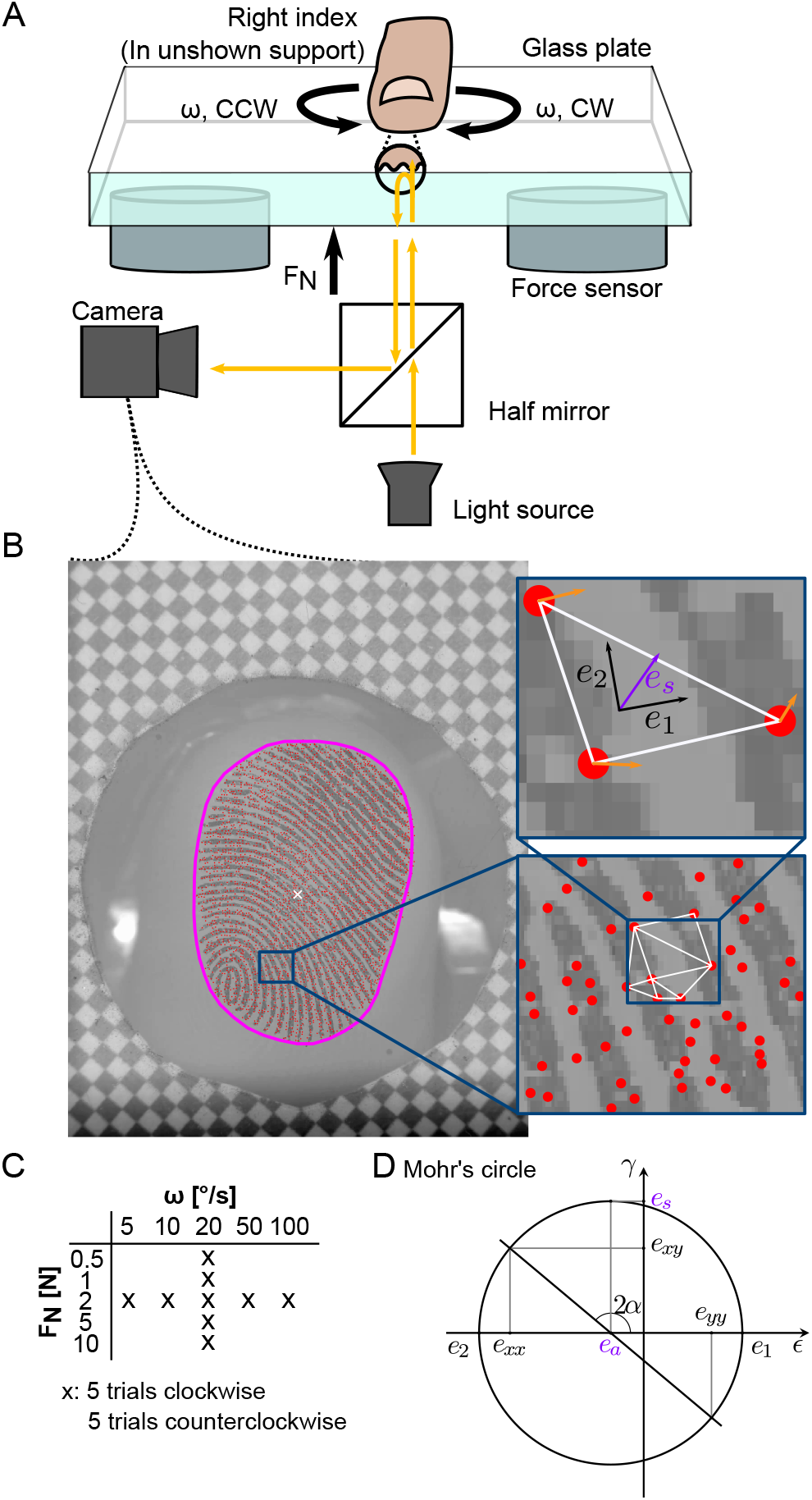
Methods. (A) Participants rested the nail of their right index finger in a support, volar side pointing down. Below, a robot controlled the position of a smooth glass plate. The normal force applied was measured by two force/torque sensors located between the robot and the plate. The finger-plate contact was illuminated and filmed through the glass plate. (B) Video frame acquired by the apparatus. The border of the contact is shown in magenta. Features of the contact were tracked (red dots) and formed the vertices of a Delaunay mesh (white triangle). Areas of skin slipping and Green-Lagrange strains were derived from the skin displacement field (orange arrows). (C) Comparisons across normal forces were made for a fixed angular velocity (20 °/s), and comparisons across angular velocities for a fixed normal force (2 N). (D) Mohr’s circle is a graphical representation of the strain tensor in multiple coordinate systems. In Mohr-circle space, the abscissa and ordinate represent respectively the normal and shear strain. Each point on the circle represents the strain tensor in a given coordinate system obtained by rotation. A rotation of α in the physical world corresponds to a rotation of 2α in Mohr-circle space. The two intersections between the circle and the abscissa represent principal strains (absence of shear strain). The radius of the circle is the maximum shear strain, obtained in the coordinate system at 45° from the principal strains (90° in Mohr-circle space). The circle centre represents the average magnitude of the two principal strains, or area invariant.

Images of the finger pad were captured using a camera (JAI-GO-5000M-PMCL) with a resolution of 2560×2048 pixels (leading to a resolution of 85 pixels/mm) and a frame rate equal to the target angular velocity in °/s (5 to 100 fps), see Fig. 1B. The finger-plate contact was illuminated by a diffuse light source and filmed through the glass plate. Forces, position, and orientation data from the robot were sampled at 1 kHz.

#### Precise measurement of tangential torque

Although the tangential torque at the centre of rotation could be estimated from the pair of sensors, the quality of the measurement was unsatisfying. Indeed the off-centered force/torque sensors will measure small tangential forces *F*_*T*_, inversely proportional to the distance of the rotation centre (T = d\**F*_*T*_), giving noisy signals. To palliate this shortcoming, the same protocol was repeated in a second experiment involving the same participants, where the platform was modified to have a single force/torque sensor directly below the finger, thereby not recording the fingerpad images. This gives a more precise measure of the tangential torque.

### Data analysis

#### Preprocessing

Force and torque data were filtered using a zero-phase Butterworth filter of order 2, with a cut-off frequency of 5 Hz.

#### Torque

To study the effect of normal force and angular velocity on the torque generated by the rotation of the plate, we extracted the magnitude of the torque during its plateau (*T*_*slip*_) as the average value on the 0.5 s window preceding a twist angle of 75° (see Fig. 2A). This angle was selected instead of 80° to avoid the deceleration phase of the plate.

**Figure 2.**
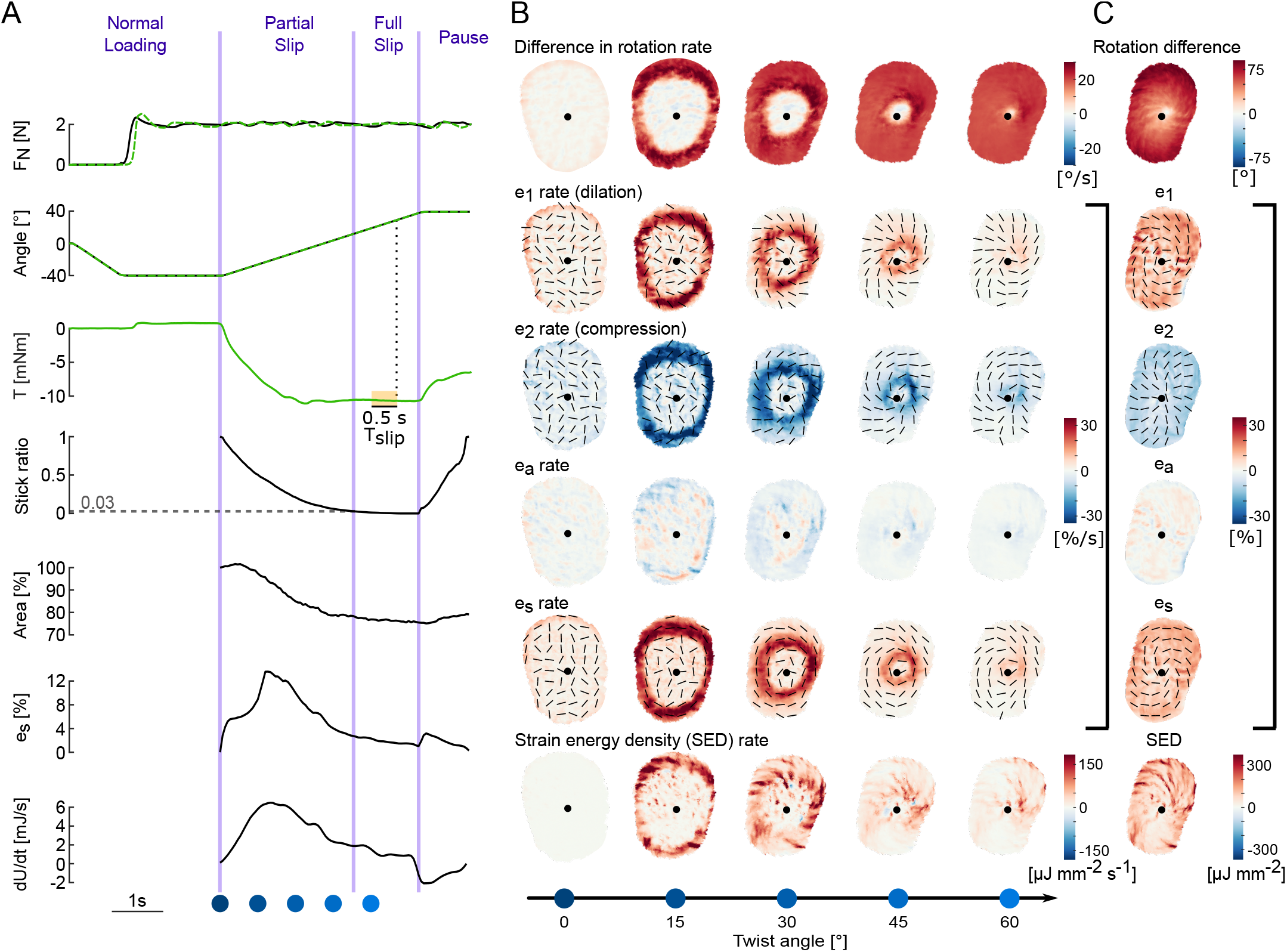
Illustrative trial. Data come both from a trial with video recording and a corresponding trial with precise torque measurements, at 2 N, 20 °/s, counter-clockwise, of participant 2. (A) Time evolution of normal force, stimulus angle, torque, stick ratio, contact area, shear strain, and strain energy for an illustrative trial. Black traces come from one trial of the experiment with video recordings, while orange traces come from a corresponding trial of the experiment measuring torque precisely with a force sensor directly under the finger. The horizontal dashed line on the stick ratio plot highlights the time and angle when full slip occurs. Blue dots correspond to the frames displayed in panel 2B. (B) Shows various descriptions of the skin strain rates in rows, with columns showing snapshots at times represented by the blue dots in Fig. 2A throughout plate rotation. The centre of rotation is marked with a black dot. **Rotation rate** of the skin is expressed in the frame of reference of the rotating plate, with increasing intensity of colour indicating a greater difference in rotation rates of skin and plate, implying that the skin is slipping. **Two principal strain rates** (*e*_1_ and *e*_2_) express the local deformation as purely dilation and compression (i.e., the local reference frame is chosen so that the shear component is zero), associated eigenvectors are indicated by black lines. **Area strain invariant** (*e*_*a*_) describes the local change in skin area. The **maximum shear rate** (*e*_*s*_) is determined by choosing the local coordinate frame such that the shear component of the local strains is maximised, at 45° from *e*_1_ and *e*_2_. Finally, the **strain energy density** is shown, which approximates the energy stored in the skin surface due to local deformations, assuming a Young’s modulus of 1 MPa and a Poisson ratio of 0.4. (C) The rightmost column shows the delta of the corresponding row variable between the first and last frame of the 80° rotation.

#### Coefficient of rotational friction

We computed the coefficient of rotational friction (μ_*rot*_), analogous to the coefficient of linear friction for each trial. The coefficient of rotational friction is defined in eq. (1):

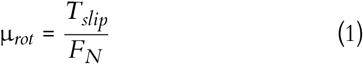

where *T*_*slip*_ and *F*_*N*_ represent the torque and the normal force during steady slip respectively.

#### Video processing

The frames of the video of the fingerpad area were filtered spatially with a Gaussian bandpass filter in the range of fingerprint ridges (0.4 mm). Optimal features were detected and tracked from frame to frame using the Lucas-Kanade-Tomasi algorithm implemented in the OpenCV toolbox (Lucas and Kanade 1981; Shi and Tomasi 1994; Bouguet 2000), and used to compute the skin displacement field. The displacement field was filtered using a convolution with a Gaussian window (N=16, σ=0.75 frames). A Delaunay triangulation of those features was computed, tessellating the contact area into triangular skin elements (Fig. 1B).

#### Apparent contact area

The apparent contact area *A*^*A*^ was segmented using a semi-automatic machine learning algorithm (see the magenta border in Fig. 1B). The size of the contact area was determined by counting the pixels inside the segmented region, and converting the result to mm^2^ using the conversion ratio of 85 px/mm given by the paper checkerboard pattern stuck on the glass and visible in every frame (Fig. 1B). The principal orientation of the contact area was approximated by the angle between the y-axis and the major axis of an ellipse fitted on the contour of the contact area following the same procedure as in Delhaye *et al*. (Delhaye, Lefèvre, and Thonnard 2014). The relative change of orientation was obtained by computing the difference between the tilt at initial contact and the tilt when fully slipping.

During the stick-to-slip transition, the contact area can be modified by two mechanisms: skin breaking or making contact, and surface deformations.

To quantify the area of skin lifting off or entering contact at the contact edges (respectively peeling or laying, see also (Lengiewicz et al. 2020) for terminology), we summed the areas of all Delaunay triangles entering or leaving contact. For a triangle to be considered in the contact area, its three vertices had to be located inside the segmented contacted area. Areas of triangles passing from inside to outside of the contact counted toward skin peeling.

To compute the influence of surface deformations on area change, we studied the frame-to-frame area ratio of all Delaunay triangles in the segmented contact area.

#### Propagation of slip

To measure the amount of skin slipping on each frame of the video, we considered that a triangular skin element had slipped if its rate of rotation differed from that of the rigid plate by more than 0.6 °/frame. The rate of rotation of a triangle was defined as the average angular displacement of the triangle vertices around its centroid. We then computed the stick ratio as the ratio of the area of skin stuck to the plate on a frame over the total area of skin in contact with the glass plate. The moment of full slip was defined as the moment when the stick ratio becomes smaller than 0.03, as slip is challenging to detect close to the rotation center where the displacement of the stimulus is almost zero.

#### Local strains

Two-dimensional Green-Lagrange strains were computed at the skin-plate interface by computing the change in the shape of each triangular element between one frame and the next. The deformations of each triangular element are therefore quantified by a strain tensor ϵ having three independent values: the normal strain along the x-axis (ϵ_*xx*_), the normal strain along the y-axis (ϵ_*yy*_) and the shear strain (ϵ_*xy*_). The complete details of the strain computation can be found in (Delhaye et al. 2016).

We found that using a single coordinate system common to all triangular elements did not yield intuitive results. While the problem at hand involves rotations making a Cartesian system unappealing, it lacks rotational symmetry (elliptical shape of the contact area and potentially the pressure distribution not being symmetrical by rotation) which would enable the use of a polar coordinate system. Instead, we expressed strains in local Cartesian coordinates, specific to each triangular skin element at each frame, and obtained by rotations based on the direction of the principal strains.

Mohr’s circle, presented in Fig. 1D, is a graphical representation of the local strain tensor in all of such rotated coordinate systems and provides a graphical intuition for the following paragraphs.

The local principal strains express the local strains in a coordinate system which removes the shear strain component. For each triangular element and between each consecutive pair of frames, it is possible to find a set of perpendicular axes where the shear strain is zero. The strains along both axes are, respectively, the maximum and minimum strains across all orthonormal coordinate systems obtainable by rotation. Those are obtained by eigenvalue decomposition of the strain tensor ϵ. The obtained strains are called principal strains and named *e*_1_ and *e*_2_ respectively. This allows us to express the local deformation of the skin as a stretch along one local axis (maximum strain, *e*_1_ > 0) and compression along the perpendicular axis (minimum strain, *e*_2_ < 0).

From the principal strains, we can compute the change in the area of each triangular element (*e*_*a*_) as:

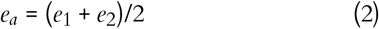

which also corresponds to the centre of Mohr’s circle (Fig. 1D). The rotation of the plate in contact with the finger will mainly cause shear strains to happen on the skin. Therefore, we computed the maximum shear strain on each triangular element as:

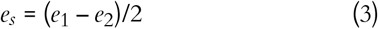

which is the radius of Mohr’s circle (Fig. 1D).

To summarise and compare shear strain across conditions, we defined |*e*_*s*_| as the median shear strain over the contact area after the full rotation.

#### Strain energy

To estimate strain energy, which is the energy stored in the skin surface as it is deformed during the rotation, Young’s modulus and Poisson ratio were chosen to match the values estimated by (Delhaye et al. 2016) which are *E* = 1 MPa and ν = 0.4. The strain energy density (SED) *u*_*d*_ was obtained for each triangle element according to:

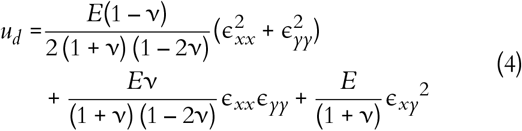

The surface SED was computed assuming homogenous deformation over a 2 mm depth, and the total energy over the contact area was evaluated following (5).

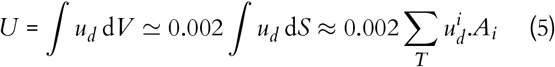

### Statistical analysis

The influence of normal force, angular velocity, and rotation direction were assessed with repeated measures analyses of variance (rANOVAs). The five repetitions of each condition were averaged. Because of the experimental design (holding force constant and varying angular velocity, or vice versa; c.f., Fig. 1C), two rANOVAs were used per dependent variable. One used normal force as an independent variable, the other used angular velocity, and rotation direction was included in both as a second independent variable. Mauchly’s test was used to test for sphericity. Greenhouse-Geisser correction was applied if sphericity was violated. Effect sizes were measured using partial eta squared 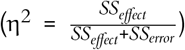. Pairwise t-tests with Bonferroni correction for multiple comparisons were used for posthoc comparisons when rANOVAs showed a significant effect of one or multiple independent variables. The significance threshold used was 0.05.

## Results

### Illustration of a single trial run

An overview of the collected data is presented in Fig. 2, in an example trial from a typical participant at the centre of the parameter space. In this example, the normal force was set to 2 N and the angular velocity was 20 °/s (see top two graphs in Fig. 2A). These typical traces show that, while the normal force was servo-controlled at a constant value, the plate was rotated a total of 80° at a constant speed for a total duration of 4 s.

As the plate started rotating, the friction between the glass and the finger skin generated a torque in the direction opposing motion (see torque *T* in Fig. 2A). This torque increased as the plate was rotated until it reached a plateau and remained stable for the rest of the rotation, at around -10 mNm in this example.

During the rotation, the transition from a fully stuck state to a fully slipping state occurs; that is, the stick ratio transitions from 1 to 0 over the course of the rotation. The progression of the slipping front can be very clearly observed in a heat map showing the difference in rotation rate between the skin and the plate (see row *Difference in rotation rate* of Fig. 2B); when the skin is slipping it is no longer rotating at the same angular velocity as the plate. The slipping front progresses toward the centre of rotation, propagating as a wavefront, and reaches the centre at the moment of full slip. Along this wavefront, we observe the most extreme values of skin deformation rate, which can be expressed either in terms of principal strain or maximum shear strain (see rows *e*_1_, *e*_2_ and *e*_*s*_ in Fig. 2B).

The principal strains, which express local deformation as purely dilation and compression, in a local coordinate system where there is no shear, are shown in the *e*_1_ *rate (dilation)* and *e*_2_ *rate (compression)* rows of Fig. 2B. Both *e*_1_ and *e*_2_ are maximal just behind the slip wavefront, with which their orientation forms a 45°angle. The skin behind the wavefront is slipping on the plate and not rotating at the same rate as the skin still stuck to the plate in front of the slip wavefront, giving rise to those strains.

We also observed in Fig. 2A (*area* subplot) that the apparent contact area decreases during the transition from fully stuck to fully slipping, similar to what is observed when the plate moves under linear tangential motion (Delhaye, Lefèvre, and Thonnard 2014). This reduction in area cannot be solely explained by compression at the surface of the skin. When looking at the area invariant *e*_*a*_ in Fig. 2B and C, there is both compression and dilation at work. Those changes in area caused by dilation and compression are distributed non-homogeneously over the contact area. When looking both at *e*_*a*_ rate and at the total *e*_*a*_ after the full rotation is complete, small wrinkles are present. As *e*_1_ and *e*_2_ (*e*_1_ and *e*_2_ lines of Fig. 2) are similar in magnitude but opposite in sign, they approximately cancel each other out, leaving the residual wrinkle pattern, as observed. As determined by visual inspection, these wrinkles do not seem to align with the fingerprint ridges. They represent small changes in area relative to the magnitude of deformation seen in the two principal strain components.

When using the local reference frame which maximises the shear component of the local deformation (at 45° of the orientations of the principal strains), we also see an intense pattern of local shear along the slip wavefront. The orientation of the principal strains *e*_1_ and *e*_2_ are at 45° to the slip wavefront, so the shear *e*_*s*_ is maximal along the radial or tangential orientations (see black orientations in Fig. 2B, for *e*_1_, *e*_2_ and *e*_*s*_). After the full rotation, *e*_*s*_ is homogeneous over the contact area (see *e*_*s*_ row in Fig. 2C). Considering both the two principal strain variables and the maximum shear variable together, we see an overall picture which is dominated by shearing of the skin along the slip wavefront. Given the small overall change in local area (*e*_*a*_), the local strains are well summarized by the local maximum shear (*e*_*s*_) and its orientation. Therefore, the subsequent heat maps (Fig. 6) only show these two elements.

The rate of change of the total energy over the contact area typically followed a bell-shaped curve, starting at 0 mJ/s Sophie du Bois de Dunilac *et al*. at movement onset, peaking at the middle of the partial slip phase and decreasing to a low, but non-zero, value at full slip (full slip does not imply homogeneous velocity field).

When the stimulus stopped rotating, elasticity of the skin made it rebound in the direction opposed to that of the stimulus. The torque reduced following this relaxation, and the contact area increased again. Negative strain energy rate is also observed, as the skin loses elastic energy. Visual inspection of the contact area traces suggests that the 1-second pause after each rotation is not sufficient to allow the skin to relax completely, as evidenced by the contact area increase appearing to still be in the early stages of an exponential relaxation pattern.

### Torque and coefficient of rotational friction

Across all the normal forces applied (0.5 to 10 N) the torque plateau *T*_*slip*_ reached values in the order of tens of mNm. As expected, the maximum torque reached during the rotation increased with the normal force, going from values around 3.91 mNm for a normal force of 0.5 N up to values around 36.70 mNm for a normal force of 10 N (Fig. 3A). The repeated measures ANOVA showed a significant effect of normal force (F=159.3, p< .001, η^2^=0.96) but not of the rotation direction (F=0.02, p=.89). Post-hoc tests between all pairs of *F*_*N*_ levels were significant.

**Figure 3.**
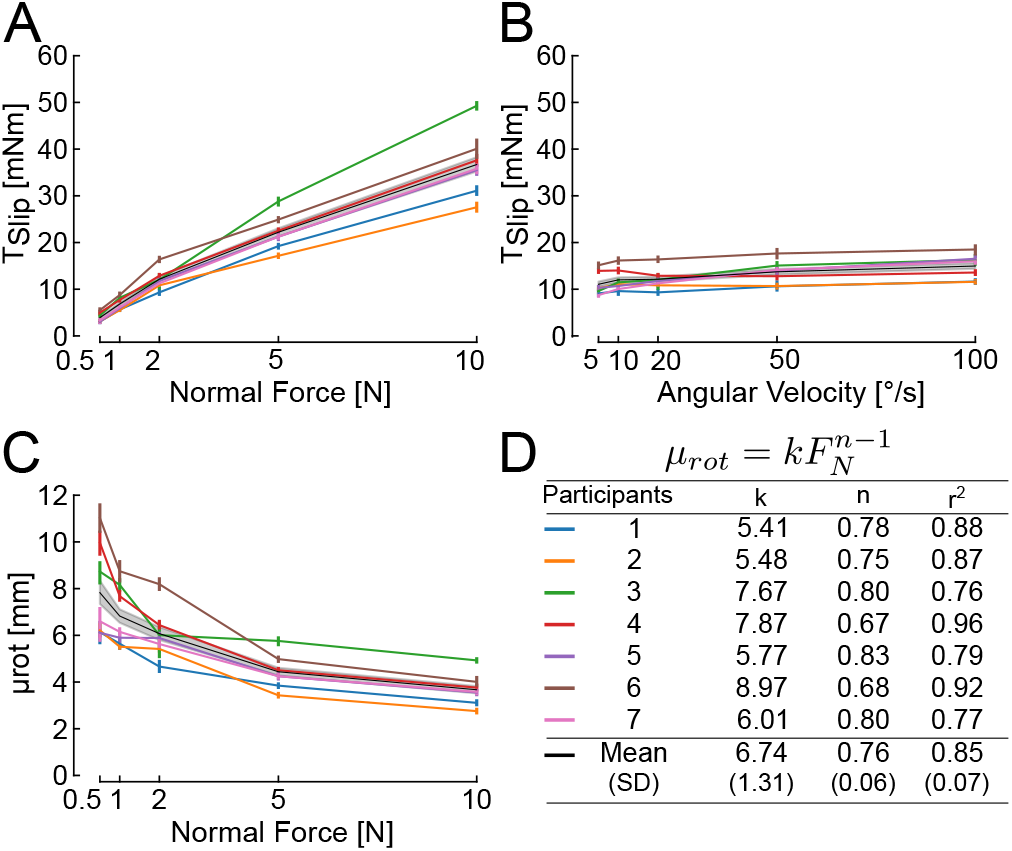
Evolution of torque plateau (T_*slip*_) with normalforce andangularvelocity, and evolution of the rotational coefficient of friction with normal force. In all panels, the black line is the mean across participants, with corresponding 95% confidence intervals in shaded grey. Individual participants are presented as light-colored lines, with colors defined in panel D. A) Maximum torque for all normal forces tested. B) Maximum torque for all angular velocities tested. C) Coefficient of rotational friction μ_*rot*_ as a function of normal force (*F*_*N*_). D) Least-square fit of the coefficient of rotational friction as a function of the normal force as the power law: 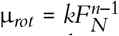.

Maximum torque also increased with angular velocity, but less than the increase due to the same relative increase of normal force (20 fold) comparing Figs. 3A and 3B. Torques increased from a value of 11.06 mNm at an angular velocity of 5 °/s to a value of 14.96 mNm at 100 °/s (Fig. 3B). The rANOVA showed there was a statistically effect of ω on *T*_*slip*_ (F=9.90, p=.017, η^2^=0.26), but the effect size was smaller than for the normal force. There was no significant effect of the rotation direction (F=0.015, p=.91). Amongst the post-hoc pairwise comparisons, only the pair 50-100 °/s reached significance.

Using the available torque and force data, we could also compute the coefficient of rotational friction μ_*rot*_ as the ratio of torque over the normal force during full slip. The coefficient of rotational friction between the plate and the participants’ finger pad varied significantly with changes in normal force, ranging from an average of 8 mm at 0.5 N of normal force to an average of 4 mm at 10 N of normal force (Fig. 3C). For comparison, the range of μ_*rot*_ reported by Kinoshita *et al*. spanned from 3.05 mm for rayon to 10.11 mm for sandpaper (Kinoshita et al. 1997), which is comparable to the range of frictions we observed. Note, in their study, no dependency of μ_*rot*_ on *F*_*N*_ was visible for rayon, suede and sandpaper.

We observed that the coefficient of rotational friction decreased non-linearly with normal force. A negative power law, 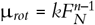, was fitted to data of individual participants to capture the relation between μ_*rot*_ and *F*_*N*_. Across participants, the parameter *k* was equal to 6.74 ± 1.31 (mean ± standard deviation) and *n* was equal to 0.76 ± 0.06 (see Fig. 3D). The coefficients of determination *r*_2_ were large, with a mean of 0.85±0.07 across participants, implying a good model fit. This result is consistent with the work on translational slip of Delhaye *et al*. who performed a similar analysis and found a value of *n* around 0.67 on glass (Delhaye et al. 2016) and Barrea *et al*. (Barrea et al. 2016) who found a value of *n* around 0.66 on Kapton.

The influence of the power law relationship between the coefficient of rotational friction and normal force (Fig. 3C) is reflected in the non-linear relationship between maximum torque and normal force in Fig. 3A. The larger coefficient of friction at lower normal forces, allows for relatively large increases in maximum torque (i.e., traction) with increases in grip force across the force ranges associated with delicate precision manipulation; i.e., 0.5 N to 2 N.

Note that the coefficient of rotational friction, as calculated here, captures two effects. One is the increasing real contact area (cumulative area of all microjunctions made between the fingerprint ridges and the glass), which increases the friction available at each local region of the fingerprint ridges. The second is that the apparent contact area increases in size with increasing normal force, with more skin coming into contact at the periphery of the contact area (further from the centre of rotation) where it can make a substantial contribution to the maximum torque reached.

### Apparent contact area

The apparent contact area after normal loading 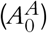 varied widely across participants, likely dependent on their finger morphology and individual biomechanics, as well as on the normal force applied. Indeed, as expected, 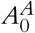 increased with the normal force (Fig. 4A), with most of the increase in initial contact area happening at low force levels (between 0.5 N and 2 N) before reaching a plateau. 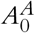 almost doubled between 0.5 N (90.23 ± ^2^ ± 23.68 mm^2^) 11.75 mm) and 10 N (182.22 across all participants. The impact of the normal force on the 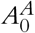 is best described by the power law, 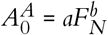, where *F*_*N*_ is the normal force, and *a* and *b* are the parameters of the fit. The mean across participants of *a* was 113.59 ± 8.48. The mean across participants of the exponent *b* was 0.21 ± 0.052. The mean of the coefficient of determination *r*^2^ was 0.93 ± 0.032, indicating a good fit. Note that participant 3 is a clear outlier from the group, achieving larger contact areas than the other six participants at 5 N and 10 N.

**Figure 4.**
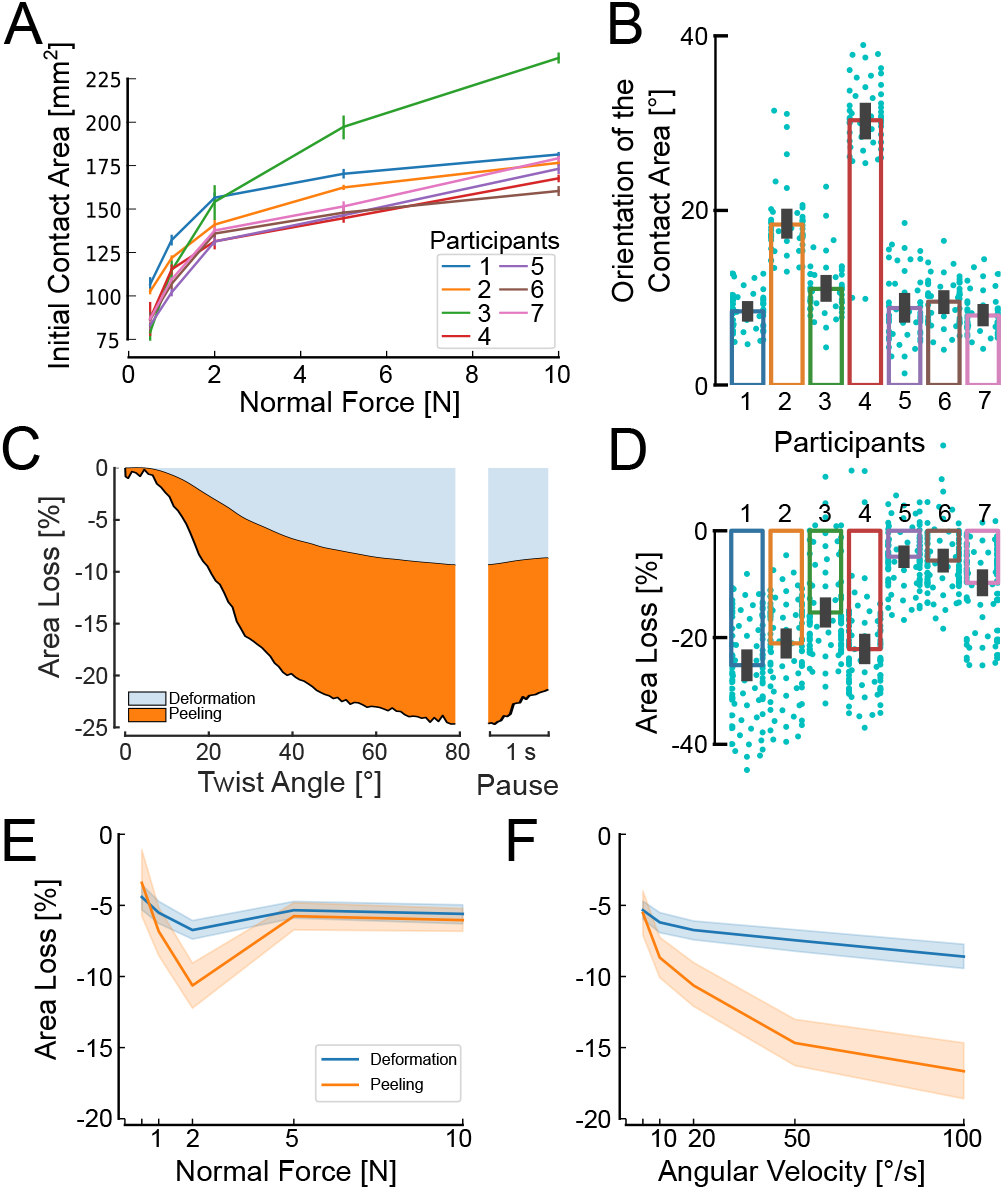
Contact Area. (A) Contact area after normal loading. (B) Change of orientation of the contact area after the 80° rotation, grouped by participant. (C) Area reduction attributable to surface deformations and peeling for an example condition and participant (average of 5 clockwise trials at 2 N and 20 °/s); (D) Relative change between initial contact area after normal loading, and contact area after the 80° rotation. (E & F) Averages across participants of the area reduction after completion of the 80° rotation as a function of normal force (E) and angular velocity (F). Error bars and shaded areas represent 95% confidence intervals.

This relationship between contact area and normal force for a human finger pad in contact with a planar surface is somewhat consistent with previous work. Delhaye *et al*. (Delhaye, Lefèvre, and Thonnard 2014) obtain power law fit parameter values ranging between 0.36 and 0.52 for *b*. However, some differences are expected, given that Delhaye *et al*. used a maximum normal force of 2 N, which is far less than the 10 N maximum normal force used in this study. Note also, the power law equation used in both this study and the paper of Delhaye *et al*. is the well-known Hertzian model of contact (with a 2/3 exponent), derived from a model of a homogeneous linear elastic sphere in contact with a planar surface (Greenwood 1985). This model is only considered valid for small contact areas relative to the size of the sphere being modelled. The shape and biomechanics of the human finger and the size of the contact area achieved relative to the size of the finger pad, violate the assumptions of this Hertzian model, and thus we expect some discrepancies between fit and experimental data.

The apparent contact area was affected by the rotation of the stimulus plate. Indeed, both its principal orientation (Fig. 4B) and its size (Fig. 4D) changed with the angular rotation of the plate. Orientation of the contact was significantly affected by normal force (F=5.45, p=.0029, η^2^ = 0.48) without clear trends being visible with no pair of *F*_*N*_ levels reaching significance in the post-hoc tests, while CW trials led to smaller changes in orientation of the contact area than CCW trials (F=10.82, p=.017, η^2^ = 0.64). The interaction direction- *F*_*N*_ did not reach significance (F=2.13, p=.15). Change in orientation of the contact area increased with angular velocity (F=24.78, p<.001, η^2^ = 0.70), and this time direction did not reach significance (F=5.23, p=.062) nor did the interaction (F=1.63, p=.25). There was a large variability across participants (Fig. 4B).

In most trials (88.83%), the size of the contact area was reduced by the rotation of the plate (see Fig. 4D). The reduction in contact area was on average 15% ± 11%. However, at low normal forces, and for only a subset of participants, an increase in contact area was observed. The reduction of the contact area was driven by two different effects: (1) peeling, i.e., the skin losing contact with the plate, and; (2) deformation, i.e., the triangular elements reducing in size. Peeling accounted for the largest component of the area reduction in most cases (Fig. 4C, E, and F).

The area reduction increased with angular velocity, with a greater increase in peeling than surface deformation (Fig. 4E). The rANOVA showed a significant influence of velocity on both the surface deformation (F=15.46, p<.001, η^2^ = 0.13) and peeling (F=33.53, p<.001, η^2^ = 0.30).

This rate dependence could imply a viscoelastic influence on the skin mechanics to cause more skin to peel from the contact area periphery at higher rates of rotation. A possible explanation is that viscoelastic effects increase the skin’s stiffness. The contact between the plate and the finger being robotically force-controlled, the plate would be retracted to restore the target force, which might contribute to skin peeling as the plate moves away from the finger.

Normal force had no significant influence on the surface deformations component of the reduction in *A*^*A*^ (F=1.32, p=.29), but affected peeling (F=2.83, p=.047). Only the pair 1-2 N reached significance in the posthoc tests, which should be interpreted with caution due to the order of presentation (see section *limitations and sources or error* of the discussion).

### Propagation of the slipping region

As expected, the periphery of the contact area started to slip first. The slip wavefront propagated towards the central region of the contact until full slip occurred (Fig. 2B).

The twist angle needed to reach full slip increased with normal force (F=14.38, p<.001, η^2^ = 0.19), see Fig. 5A, but only the pair 0.5-10 N reached significance in the posthoc tests. Full slip was reached later in CCW than in CW trials (F=8.55, p=.027, η^2^ = 0.0065), and there was no significant interaction between rotation direction and *F*_*N*_ (F=1.34, p=.28).

**Figure 5.**
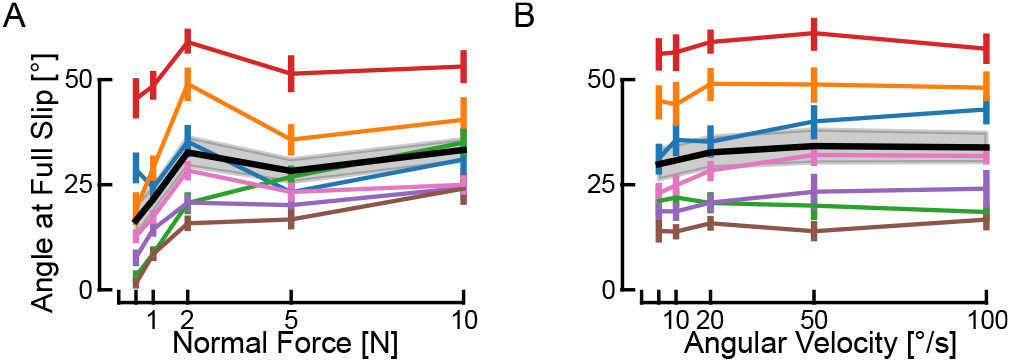
Reaching full slip. Averages (black lines) with 95% confidence intervals (grey shaded areas) across participants of the angle to reach full slip as a function of normal force (A) and as a function of angular velocity (B). Participant averages are indicated as colored lines, with 95% confidence intervals.

**Figure 6.**
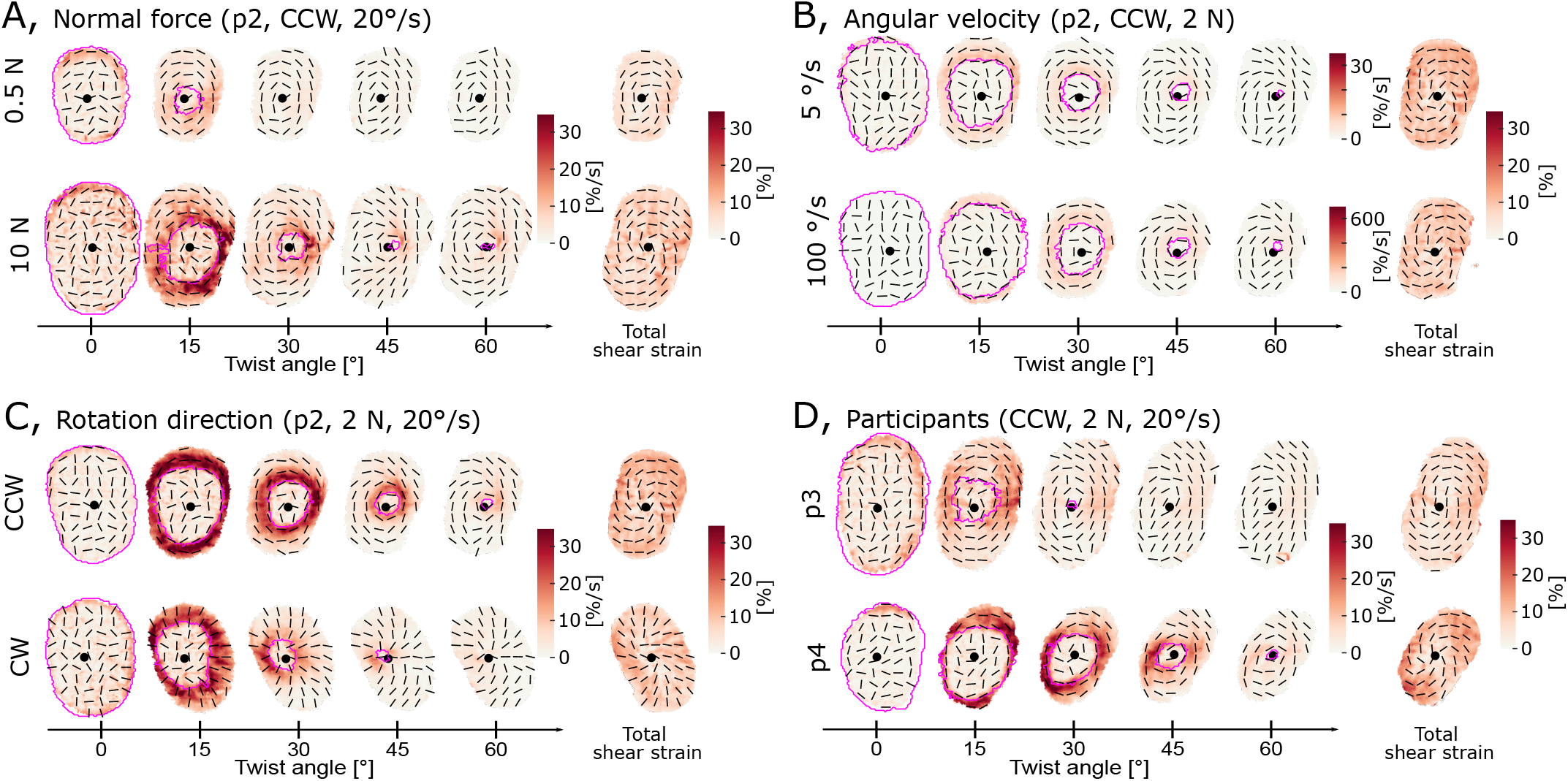
Illustrative maximum shear across conditions. The centre of rotation is indicated by a black dot, the slip wavefront by a magenta contour, and the maximum shear strain’s orientation by black lines. (A) *Comparison of different normal forces, at 20 °/s*. Comparison of maximum shear across finger pad skin for a normal force of 0.5 N versus 10 N, both for a counterclockwise trial with a rotation rate of 20 °/s (Participant 2). The heat map across each finger pad contact area illustrates the maximum shear rate at each skin location. The last column shows the total of the maximum shear for the full 80° rotation. As expected, the contact area is larger for the larger normal force. More slip happens earlier at the smaller normal force. (B) *Comparison of different angular velocities, at 2 N*. Comparison of maximum shear across finger pad skin for an angular velocity of 5 °/s versus 100 °/s, both for a counterclockwise trial with a normal force of 2 N (Participant 2). Since the rates of rotation differ by a factor of 20, for visualisation purposes, we make the scale bar ranges also differ by this factor. Despite the large difference in the rotation rate between the two trials, there is very little difference in the spatial and temporal patterns of maximum shear rate, aside from their differing magnitudes. (C) *Comparison of rotation direction, at 2 N, 20 °/s for participant 2*. (D) *Comparison of two participants, at 2 N, 20 °/s*. An illustrative comparison of the time course of single trials taken from two different participants. We observe that, while the overall pattern of the time series is generally similar across these two (and all) participants, the timing of events does vary between participants; for example, Participant 3 (p3) experiences a lot of slip early in the rotation, whereas Participant 4 (p4) achieves the same amount of slip at a later time (and larger angle of rotation).

Increasing the angular velocity had no practical impact on the rotation angle at full slip (Fig. 5B), which conversely means that it linearly shortened the time available to the participant should they have needed to react before full slip occurred. Although the rANOVA found a significant effect of angular velocity (F=6.01, p=.0017, η^2^ = 0.50), no pairs of ω showed significant differences after Bonferroni correction. There were no significant effects of direction (F=2.70, p=.15), nor interaction (F=1.89, p=.14).

### Local strains

The central portion of the contact area sticks to the plate and moves as a rigid-body rotation, while skin in the periphery slips, slows down, then stops. Surface strains thus arise behind the slip wavefront.

Aggregating local skin deformations across the finger pad, for all subjects, and over the time course of all trials into summary heat maps risks obfuscating the rich patterns that emerge throughout the slipping duration. We observe very similar patterns across all participants, which differ mostly in the times at which different amounts of slip have occurred, the contact area size after normal force loading, the amount of rotation of the contact area throughout the trial, or the intensity of the deformations experienced on the slip wavefront as it propagates. Therefore, we provide some representative examples in Fig. 6 and comparisons to help illustrate these patterns to the reader. In addition, we used the median maximum shear over the contact area |*e*_*s*_| to allow quantitative comparisons between conditions.

#### Principal strains and maximum shear

In addition to increasing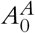 and delaying full slip, increasing the *F*_*N*_ also increased local strain rates (Fig. 6A). This is also visible as a significant influence of *F*_*N*_ on |*e*_*s*_| (F=24.53, p<.001, η^2^ = 0.26).

Angular velocity greatly influenced the strain rates expressed in %/s (pay attention to the different scales in the two rows of Fig. 6B). However, it did not have a strong effect on strain rates in %/°, meaning that the total strain after the full rotation was completed did not strongly differ between levels of ω. Despite reaching statistical significance, ω only showed a small effect size on |*e*_*s*_| (F=3.55, p=.021, η^2^ = 0.01).

CCW trials resulted in slightly higher shear strains than CW trials, and mirrored shear patterns (Fig. 6C). Although it reached significance in both rANOVAs testing the effect of *F*_*N*_ and ω on |*e*_*s*_|, the effect size was small in both cases (*F*_*N*_ : p=.0042, η^2^ = 0.02 and ω: p=.015, η^2^ = 0.01).

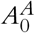, twist angle at full slip, local strain rates, and total strains all varied across participants, as illustrated in Fig. 6D.

#### Strain energy

The maximum strain energy rate 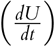 and the total strain energy (*U*) (F=22.39, p<.001, η^2^=0.79) both increased with increasing normal force (Fig. 7A). This is expected for two reasons. Firstly, a larger normal force will result in a larger *A*^*A*^, and so the opportunity for more skin to experience deformation (see Fig. 4). Secondly, a larger normal force will provide greater traction at any local region of skin, and so the propensity to generate more extreme deformations of the skin; this is evidenced by increasing maximum torque with increasing normal force (see Fig. 3A).

**Figure 7.**
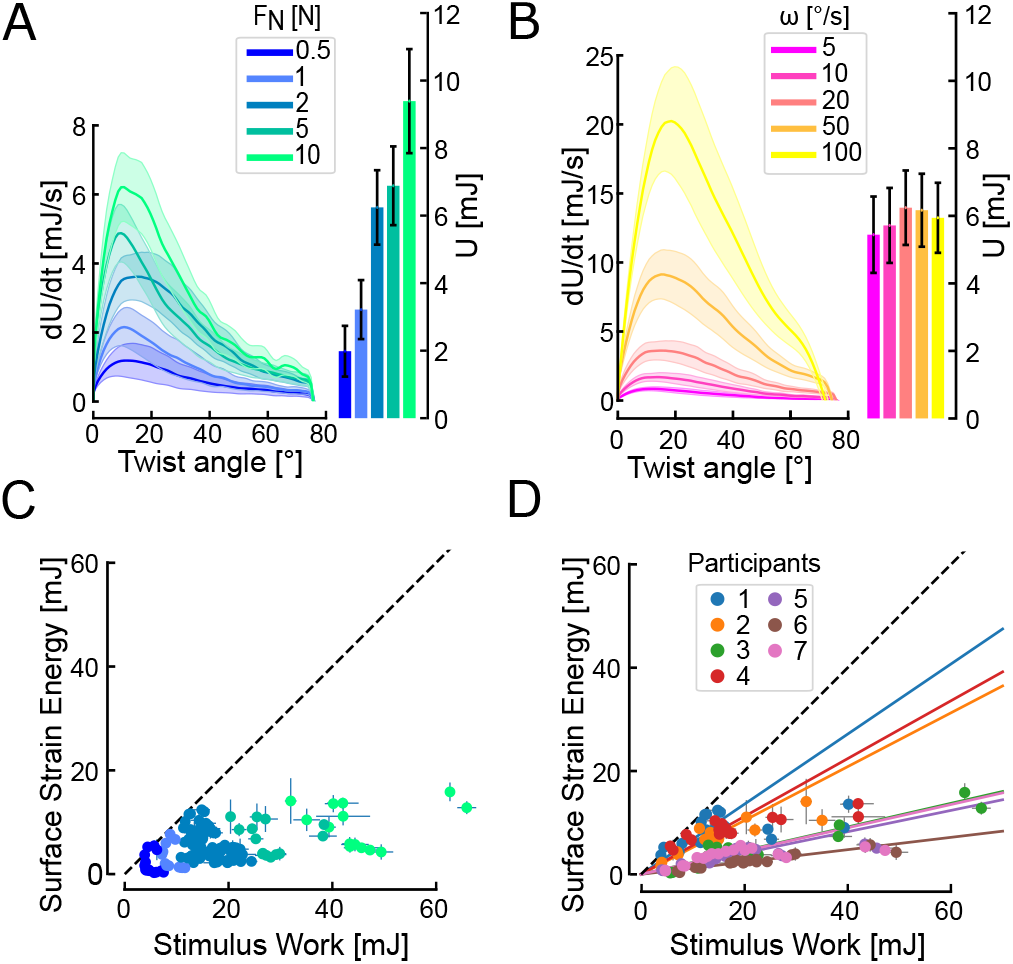
Evolution of strain energy. The influence of normal force (A) and angular velocity (B) on the evolution of the total strain energy rate 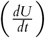 with respect to stimulus displacement (line graphs) and final total strain energy (bar charts). Each trace and box shows the mean across subjects. Shaded areas and bars show standard error. Note that in (B), the higher ω, the less time the rotation lasts, which is why despite having higher energy rates fast angular velocities do not lead to higher time integrals. *Surface strain energy versus external work*. The work provided by the stimulus is the upper limit on surface deformation energy (dashed line). Only the colour code changes between panels C and D. (C) Normal force increases the stimulus work as well as the surface deformation energy. (D) When colour coding by participants, it becomes apparent that the proportion of available work converted into surface deformation decreases with increasing normal force. Linear fits constrained to pass through the origin do not fit the data very well for participants with large deformation.

Although the maximum strain energy rate 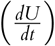 increased with angular velocity, the total strain energy (*U*) did not (F=1.46, p=.24) (Fig. 7B, right). Indeed, for larger angular velocities, the strain energy peak increased, whereas the duration of the rotation was reduced in proportion, leading to similar total strain energy.

The surface strain energy was compared to the external work applied to the finger by the stimulus (i.e., integral of torque-angle product over time). The proportion of work used to deform the skin reduced with normal force for the participants undergoing the most surface deformation (note how the higher work points deviate from the linear fits for participants 1, 2, and 4 in Fig. 7D).

It can be noted that as normal force increases, more external work is done by the stimulus and more strain energy deforms the skin. However, as the normal force increased, a smaller fraction of the available work was converted into surface deformation (Fig. 7C). Excess energy could be converted into deformation of the skin bulk, and some into heat. It should be noted that differences across subjects are most probably related to differences in skin properties (e.g. stiffness), which are set constant in the calculation.

## Discussion

We studied the effects of a torsional load on the finger pad of human participants by pressing a glass plate against their skin at a robotically-controlled normal force, then rotating it at a constant angular velocity. As the stimulus plate rotated, we observed partial slippage of the skin in contact with the plate. The periphery of the contact area started slipping first, while the region closer to the centre of rotation stuck to the stimulus plate and moved as a rigid body. While this incipient slip pattern has been well-documented for translational movements, this is the first study to comprehensively describe the finger pad skin mechanics under torsional stimulation. We observed that at the slip wavefront, an annulus of strains dominated by shear is present. While there are differences in the magnitudes and timeline of skin deformations between participants, trials generally follow a similar pattern for all participants across all experimental conditions.

### Effect of force and velocity

We tested a wide (20-fold) range of angular velocities and normal forces, using values relevant to both object manipulation and tactile exploration. We observed a significant increase in torque with both normal force and angular velocity. The effect of normal force was expected from the Coulomb model of friction (*F*_*T*_ = μ*F*_*N*_). Indeed, an increasing normal force deforms the finger pad skin, thus increasing both the apparent contact area *A*^*A*^ and also the real contact area (number and strength of atomic/molecular bonds giving rise to friction forces, which is a subset of the visible fingerprint ridge contact area (Sahli et al. 2018)). With the increase in contact area, the amount of traction available will increase and lead to a higher torque reached at full slip. However, the fact that the increase in torque with *F*_*N*_ is sub-linear (also evidenced by the decrease of μ_*rot*_ with normal force), highlights the limited validity of the Coulomb model for the finger pad.

Maximum torque is also affected by angular velocity, with small increases in maximum torque achieved for an increase in angular velocity (see Fig. Fig. 3B). This relationship would be somewhat expected, given the skin’s known viscoelastic properties; one would expect skin stiffness to be dependent on the rate of change of internal forces within the skin as it is deformed, and so make the skin appear stiffer when deformed more rapidly at larger angular velocities.

### Comparison between rotating and translating stimuli

The present study extends the work done using rectilinear stimuli (Delhaye, Lefèvre, and Thonnard 2014; Delhaye et al. 2016). In both cases, partial slippage of the contact area was observed, starting at its periphery. The skin at the centre of the contact area slips last, while surface strains are present at the slip wavefront. This similarity between slips under rotation and translation was expected due to the curvature of the finger pad resulting in smaller traction at the periphery of the contact area, which is where slips first occur when the contact is challenged by shear forces.

Despite this similarity, there are differences in strain patterns between rectilinear and torsional motion. In rectilinear motion, there is compression at the leading edge and dilation at the trailing edge, accompanied by net change in area locally. In rotation, compressive and dilative principal strains, which are similar in magnitude, give rise to a strong shearing effect within the incipient slip wavefront annulus, but only small area changes. Although their patterns differ, strains in both types of motion contain information regarding the safety of the contact.

The present study proposes maximum shear as a tool to study surface skin deformations. We showed that it is largely present in rotations. It has the advantage of being an invariant of the strain tensor, bypassing the need to define a fixed coordinate system, which is not a trivial task when studying complex motions, be it passive rotations or combinations of translations and rotations encountered during natural object manipulation.

### Limitations and sources of error

#### Linking torque and deformations

Because of the noisy recording of torque during experiment 1, torque had to be acquired during a second experiment. This limits the possibility of precisely linking time points in the torque evolution with skin deformations and contact area reductions. Indeed, as the experiments were conducted on separate days, parameters such as the moisture of the participant’s finger could have impacted the friction. However, given that both experiments were performed with the same participants and the same experimental equipment, very similar variations in the behavior of their fingers were achieved, hence allowing us to qualitatively compare the results of experiment 1 and experiment 2. One possible way to circumvent this issue would be to use a ring-shaped force sensor, rather than a disc, allowing the finger pad to be imaged through the hole in the ring-shaped sensor.

#### Effect of the order of presentation

Data points at 2 N do not always fit the general trends, such as for area loss (Fig. 4E), the twist angle at full slip (Fig. 5A), or the strain energy (Fig. 7A). The order of presentation of the trials at different normal forces was not randomized, as the focus of the camera needed to be adjusted for each normal force. The trials were presented in blocks of similar normal force in the following order: 0.5 N, 1 N, 5 N, 10 N, and 2 N. Two effects are at play: change of normal force and total experiment time. The latter might have an influence on finger characteristics (such as moisture). In hindsight, this effect could have been mitigated by changing the presentation order across participants.

### Potential implications for object manipulation

Trial-to-trial adaptation of grip force is slower in the presence of torque, possibly explained by differences in cutaneous feedback received (Crevecoeur et al. 2011). In the present study, we confirm that local deformation patterns are qualitatively different under torsion than translation. In addition for similar minimum required grip forces, there might be more partial slips in the presence of torque, as at the periphery of the contact area there is both a decrease in pressure and an increase in constraints. Devising an active manipulation experimental paradigm comprising both torque and imaging of cutaneous deformations could help investigate whether the amount of partial slips and/or the nature of the strains (shear vs. area change) created affect grip force adaptation.

### Variability between participants

An important variation in the biomechanical response of the skin was observed across subjects. A typical example of such variation concerns the angle needed to reach full slip: Fig. 5 shows that there is almost a four-fold difference across participants in the twist angle required to reach full slip (for example, at 2 N and 20 °/s, subject averages ranged from 15.81° to 58.93°). The consequence of this, given that the plate rotates at a constant angular velocity, is that the time to full slip varies between participants. In addition, Fig. 6D provides an illustrative comparison of slip propagation for Participant 3 (p3) and Participant 4 (p4). It shows the difference in the timing of the strains’ evolution and highlights the difference in strain magnitude experienced by participants. This wide range of responses of the skin to a single stimulus was also pointed out by Wang and Hayward (Wang and Hayward 2007) and illustrates the need to pay attention to biomechanical variables when searching for the causes of sensory afferent responses, instead of focusing solely on gross mechanical stimulation parameters, such as the time course of tangential force and torque, or velocity, or stimulus curvature (Birznieks et al. 2001; Birznieks et al. 2010; Loutit et al. 2022; Goodwin, Browning, and Wheat 1995). Although some afferent types (slowly adapting) are sensitive to the resulting force vector, others (fast adapting type I) might correlate better with local deformations (Delhaye et al. 2021). It is also an argument against the use of standardised fingers when reporting data from multiple participants. Two afferents reported at the same location on a standardised finger might be subjected to very different strains, even if the external stimulus was well controlled. Ultimately, afferents respond to local events that trigger their end-organs and ignoring the variation in skin dynamics between subjects might blur subsequent analyses. Linking observed events as close as possible to the receptive fields of afferent, such as surface skin deformation, can mitigate this effect. In addition, statistical tools, such as linear mixed effect models, which take into account the nested structure of data, could prove useful when relating microneurography recordings with mechanical variables (Yu et al. 2022). It is likely that other factors, such as the orientation and depth of an afferent in the skin, will also lead to different responses from afferents of the same type. Hence, avoiding the use of standardised fingers in the analysis of afferent response will improve the understanding of the differences in afferent behavior but will not totally explain these differences.

### Potential links with mechanoreceptor firing

For rectilinear motion, surface strains have been linked to afferent firing (Delhaye et al. 2021), and it is likely that a similar link exists in rotation that further microneurography studies would highlight. Furthermore, microneurography studies with a broader variety of mechanical stimuli might help pinpoint which aspects of the skin deformation cause afferents to fire. For example, an experiment with rotation, as in this paper, would not allow for the determination of whether compression, dilation, or shear, which are all simultaneously present on the slip wavefront, are primarily responsible for afferent firing. Whereas as highlighted above, translating stimuli show more area changes and less shearing than rotating stimuli, and thus it might be possible to use a paradigm involving both stimuli types to disentangle the sensitivity of the afferents under investigation to specific strains.

In the current study, we highlighted skin peeling at the periphery of the contact area, and some mechanoreceptors might be sensitive to that. SA-I afferents are typically described as sensitive to indentation (Handler and Ginty 2021) and might respond to the skin losing contact with the plate. A limitation, however, is that with experiments measuring surface deformation only, we do not have information about the deformation of the bulk of the finger pad in which the sensory afferents are embedded, or other locations outside the contact area such as the nail bed. Alternative techniques to study skin deformation include optical coherence tomography (OCT) (Lee et al. 2020), multi-camera systems (Kao, Xu, and Gerling 2022), or computational modeling such as finite element modeling (Dandekar, Raju, and Srinivasan 2003; Wang et al. 2016).

## Conclusion

In this work, we showed the complex skin deformations at the interface between the human finger pad and a glass plate under a torsional load. We highlighted the important shear effects at the periphery of the contact, between the stuck and slipping parts of the skin. These mechanics are important as they play a significant role in generating the tactile information used during manipulation. We further show that the mechanics of the skin vary widely across participants, hence highlighting the importance of accounting for such variability when ensembling the results of neurological recordings from multiple participants onto a standard finger. In future work, we would like to combine microneurographic recordings with this imaging approach and a novel paradigm involving both translation and rotation, to better understand how different strain types (dilation, compression, and shear) contribute to the responses of different afferent types, and hence our ability to sense grip security. We will also consider other imaging modalities which allow us to measure skin deformation outside of the contact area between the skin and the plate; such as at the sides or in the bulk of the finger.

